# Genetic interactions contribute less than additive effects to quantitative trait variation in yeast

**DOI:** 10.1101/019513

**Authors:** Joshua S. Bloom, Iulia Kotenko, Meru J. Sadhu, Sebastian Treusch, Frank W. Albert, Leonid Kruglyak

## Abstract

Genetic mapping studies of quantitative traits typically focus on detecting loci that contribute additively to trait variation. Genetic interactions are often proposed as a contributing factor to trait variation, but the relative contribution of interactions to trait variation is a subject of debate. Here, we use a very large cross between two yeast strains to accurately estimate the fraction of phenotypic variance due to pairwise QTL-QTL interactions for 20 quantitative traits. We find that this fraction is 9% on average, substantially less than the contribution of additive QTL (43%). Statistically significant QTL-QTL pairs typically have small individual effect sizes, but collectively explain 40% of the pairwise interaction variance. We show that pairwise interaction variance is largely explained by pairs of loci at least one of which has a significant additive effect. These results refine our understanding of the genetic architecture of quantitative traits and help guide future mapping studies.

## Introduction

Genetic interactions arise when the joint effect of alleles at two or more loci on a phenotype departs from simply adding up the effects of the alleles at each locus. Many examples of such interactions are known, but the relative contribution of interactions to trait variation is a subject of debate^1–5^. We previously generated a panel of 1,008 recombinant offspring (“segregants”) from a cross between two strains of yeast: a widely used laboratory strain (BY) and an isolate from a vineyard (RM)^6^. Using this panel, we estimated the contribution of additive genetic factors to phenotypic variation (narrow-sense or additive heritability) for 46 traits and resolved nearly all of this contribution (on average 87%) to specific genome-wide-significant quantitative trait loci (QTL). The repeatability of trait values across replicate measurements for each segregant provided an upper bound for the total contribution of genetic factors to phenotypic variation (broad-sense or full heritability). We used the difference between trait repeatability and the additive heritability as an estimate of the contribution of genetic interactions to trait variation. Because trait repeatability can include sources of variation other than gene-gene interactions, this approach can overestimate the contribution of such interactions. Further, with 1008 segregants, we were able to detect only a small number of significant QTL-QTL interactions that, in aggregate, explained little of the estimated interaction variance.

Here, we address these limitations by studying an expanded panel of 4,390 segregants obtained from the same cross. We genotyped these segregants at 28,820 unique variant sites and phenotyped them for 20 end-point growth traits with at least two replicates. The larger sample size permits us to directly and accurately quantify pairwise interaction variance, instead of relying on the difference between trait repeatability and the additive heritability. It also greatly increases the power to detect both additive QTL and QTL-QTL interactions (Supplementary Fig. 1). For example, we have 90% power to detect an additive QTL that explains 0.5% of phenotypic variance, and 90% power to detect a QTL-QTL interaction that explains of 0.8% of phenotypic variance (Methods). Further, the expanded panel substantially improves fine mapping of loci.

We detected nearly 800 significant additive QTL. We were able to refine the location of the QTL explaining at least 1% of trait variance to approximately 10 kb, and we resolved 31 QTL to single genes. We also detected over 200 significant QTL-QTL interactions; in most cases, one or both of the loci also had significant additive effects. For most traits studied, we detected one or a few additive QTL of large effect, plus many QTL and QTL-QTL interactions of small effect. We find that the contribution of QTL-QTL interactions to phenotypic variance is typically less than a quarter of the contribution of additive effects. These results provide a picture of the genetic contributions to quantitative traits at an unprecedented resolution.

## Results

### Partitioning trait variance

We used a linear mixed model with additive, pairwise interaction, and residual strain repeatability terms to quantify these components of trait variation^7^. The additive and interaction genetic contributions are estimated based on the realized relatedness^8^,^9^ of all pairs of segregants, as measured from the dense genotype data. This approach allows us to separate the contribution of gene-gene interactions from other genetic and non-genetic sources of variation that can contribute to trait repeatability^7^. We used simulations (Methods) to demonstrate that the model can accurately estimate the contributions of additive QTL and QTL-QTL interactions to trait variation over an extensive range of genetic architectures (Supplementary Fig. 2 and Supplementary Data 1).

Across the 20 traits, additive genetic variance ranged from 8.6% to 70.4% of phenotypic variance, with a median of 43.3%. Interaction genetic variance ranged from 2.2% to 21.2% of phenotypic variance, with a median of 9.2%. These measures provide genome-wide estimates for the aggregate effects of all additive and all pairwise interaction effects, respectively. The contribution of pairwise interactions to trait variance is typically less than a quarter of the contribution of additive effects, and does not exceed half the contribution of additive effects for any trait studied here. The remaining strain repeatability variance ranged from 0.05% to 21.4%, with a median of 8.8% (Fig. 1). Three-way interactions may account for some of the remaining effect of strain, but are unlikely to explain most of this remaining variance for most traits (Supplementary Data 2). This leaves higher-order interactions, other effects of strain, or experimental effects confounded with strain as the potential sources of the remaining strain repeatability variance.

**Figure 1.**
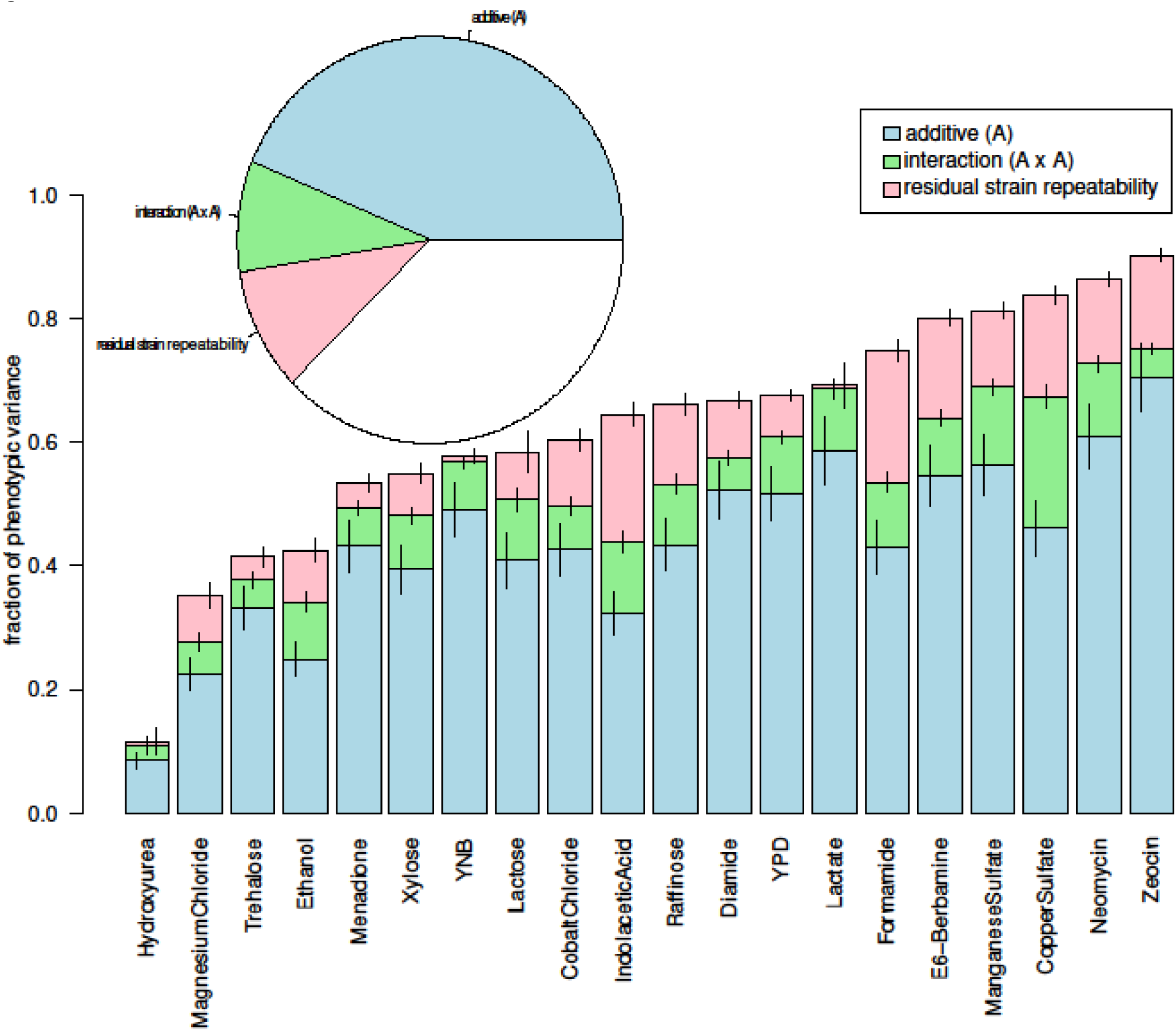
Contributions to trait variation. Stacked bar plots of a variance component analysis for each trait are shown. The variance component model included terms for additive genetic variance (blue), two-way interaction variance (green), residual strain repeatability (pink), and residual error (not shown). Error bars show +/− s.e. Inset, the average of the variance components across traits. Additive genetic effects, two-way interactions, and residual repeatability account for 43%, 9%, and 10% of phenotypic variance, respectively.

### Mapping additive QTL

Next, we sought to identify the individual genomic regions underlying these genome-wide estimates. We used a forward-search QTL mapping approach that controls for other QTL^10^ (Methods) to detect 797 genome-wide-significant additive QTL, with a median of 42.5 per trait (range 17–56). We calculated the variance captured by these detected QTL with a random effect model that uses a genetic relationship matrix (GRM) constructed only from genotypes at the peak markers for each significant additive QTL. These loci captured a median of 92% of the additive genetic variance (Fig. 2a). The number of detected QTL per trait increased approximately four-fold relative to that in our previous study of a subset of 1008 segregants from this panel^6^, but the variance captured by significant QTL only increased by 5%, because most detected loci generally have very small effect sizes (median effect size of 0.38%) (Fig. 4). These observations suggest that many additional undetected loci for these traits likely exist in this cross, but that their individual and collective effects are very small. The increased panel size also increases mapping resolution. The 180 loci that explain 1% or more of phenotypic variance have a median 95% confidence interval of 10.3kb, compared to 31.2kb with 1,008 segregants; these confidence intervals span approximately 5 genes in the yeast genome. In 31 cases, QTL could be refined to a single gene (Supplementary Data 3).

**Figure 2.**
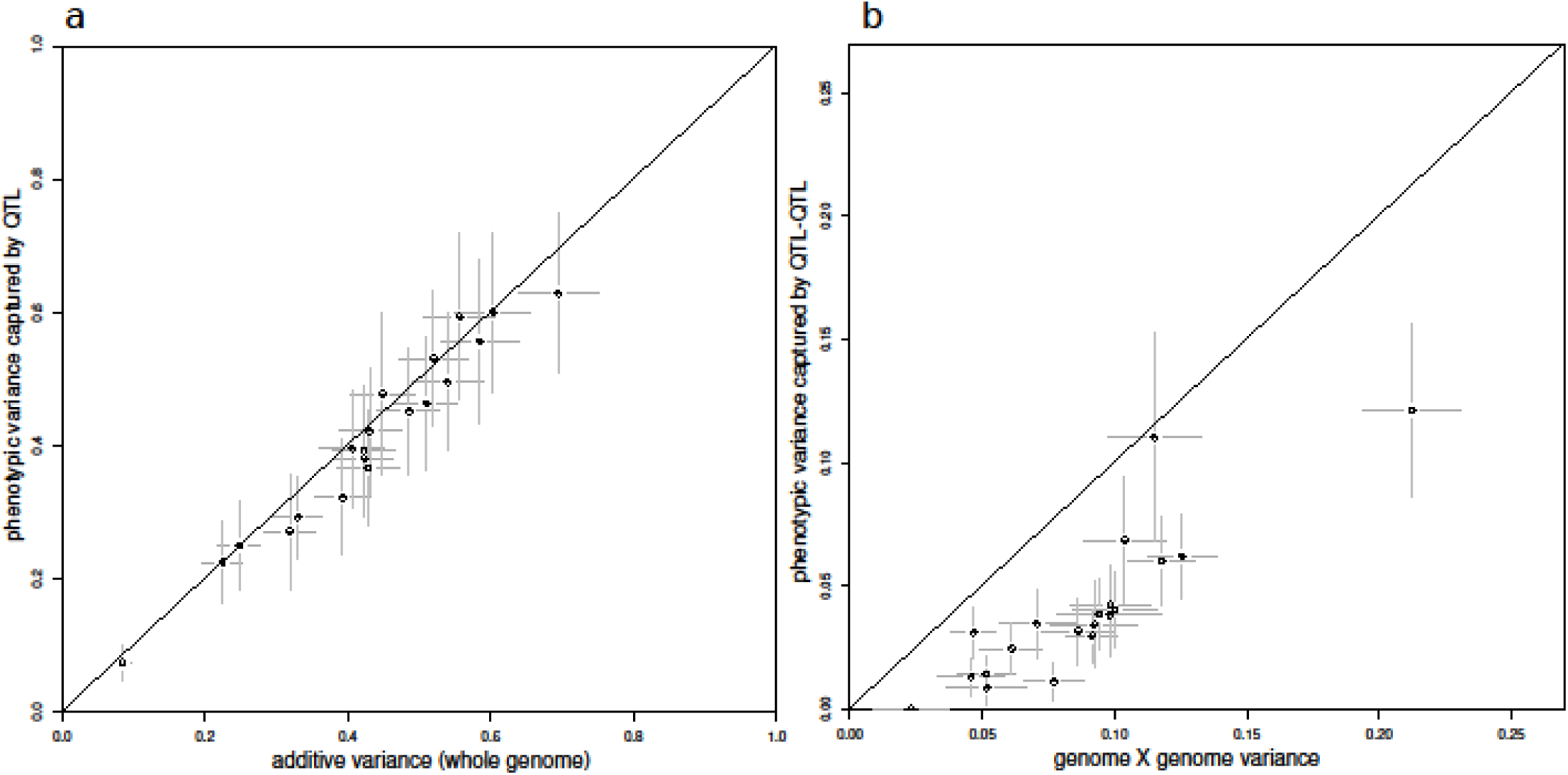
Additive and interaction variance captured by detected loci. a) Total variance captured by detected QTL for each trait is plotted against the whole-genome estimate of additive genetic variance. Error bars show +/− s.e. The diagonal line represents (variance captured by detected QTL = additive genetic variance) and is shown as a visual guide. b) Total variance captured by detected QTL-QTL interactions from the marginal scan for each trait is plotted against the whole-genome estimate of interaction variance. Error bars show +/− s.e. The diagonal line represents (variance captured by detected QTL-QTL interactions = interaction genetic variance) and is shown as a visual guide.

### Partitioning interaction variance

Detection of additive QTL that account for nearly all of the additive genetic variance allowed us to further partition the variance contributed by QTL-QTL interactions (Methods). Briefly, we compared estimates of interaction variance captured by pairs of markers selected by three different criteria: all pairs of markers across the genome, the subset of pairs in which one marker is the peak of an additive QTL, and the subset where both markers are additive QTL peaks. As noted above, across the traits examined, the amount of phenotypic variance captured by interactions between all marker pairs had a median of 9.2%. The amount of phenotypic variance captured by interactions between significant additive QTL and the rest of the genome had approximately the same median (9.4%), whereas it dropped to 4.5% for interactions only between significant additive QTL (Fig. 3 and Supplementary Fig. 3). These results suggest that in most pairwise interactions, at least one of the loci has a significant additive effect, as can be confirmed by directly mapping QTL-QTL interactions (see below).

**Figure 3.**
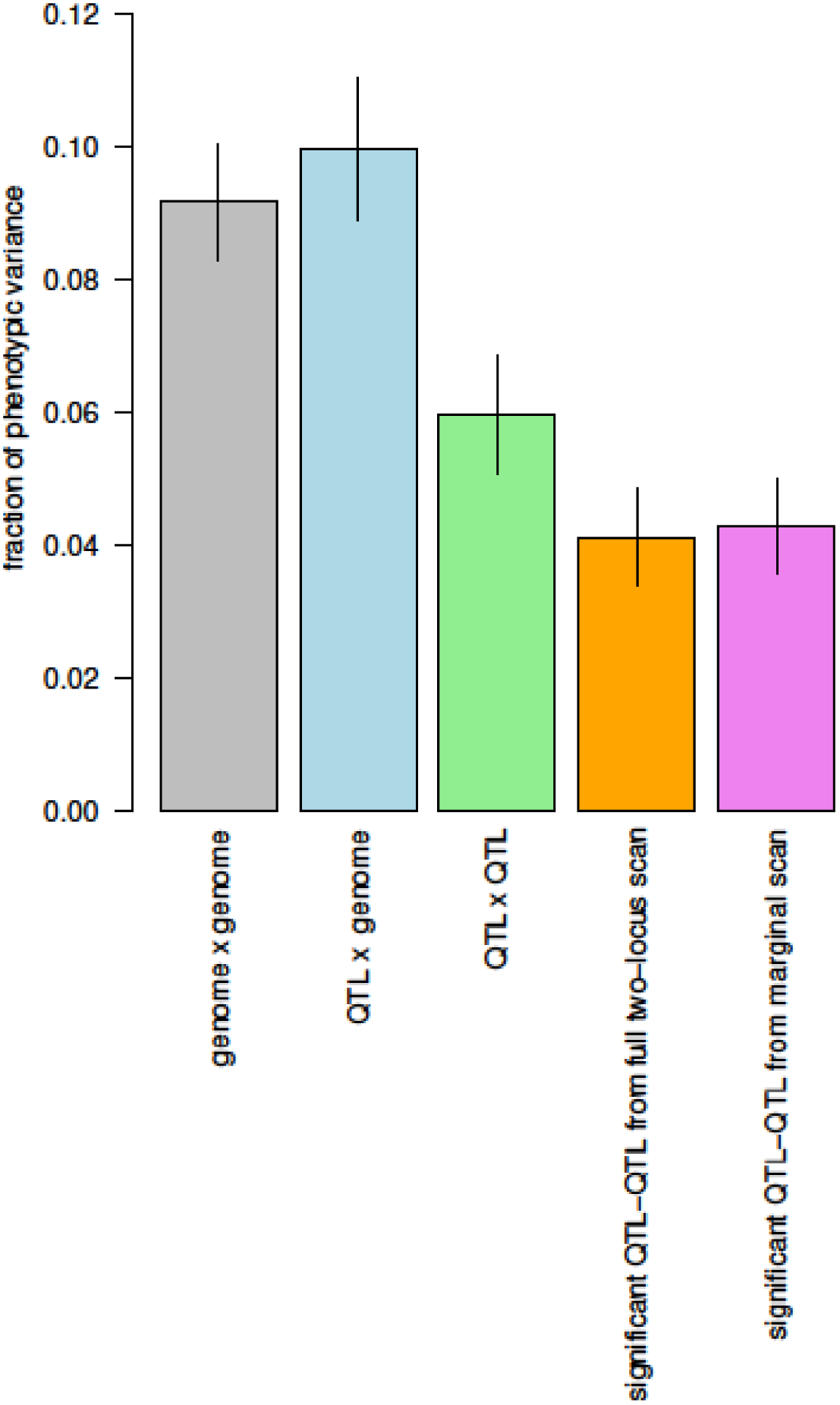
Phenotype variance captured by different variance component models of two-way interactions. The average fraction of phenotypic variance captured by different variance component models of two-way interactions across traits. The bar heights represent variance estimated with all markers (genome x genome) (grey), significant additive QTL by all markers (QTL x genome) (blue), additive QTL by additive QTL (QTL x QTL) (green), significant QTL-QTL detected from the marginal scan (orange), and significant QTL-QTL from the exhaustive two-dimensional scan (purple). Error bars show +/− s.e.

### Mapping QTL-QTL interactions

We detected specific genome-wide significant QTL-QTL interactions for each trait using a statistically powerful approach that takes into account all the additive genetic variance (Methods). One can test for interactions either between all pairs of markers (full scan), or only between pairs where one marker corresponds to a significant additive QTL (marginal scan). In principle, the former can detect a wider range of interactions, but the latter can have higher power due to a reduced search space. Here, the two approaches yielded similar results, detecting 205 and 266 QTL-QTL interactions, respectively, at an FDR of 10%, with 172 interactions detected by both approaches. In the full scan, 153 of the QTL-QTL interactions correspond to cases where both interacting loci are also significant additive QTL, 36 correspond to cases where one of the loci is a significant additive QTL, and only 16 correspond to cases where neither locus is a significant additive QTL (Supplementary Fig. 4 and Supplementary Data 4). The interactions detected in the full and marginal scans captured a median of 3.2% and 3.4% of phenotypic variance, respectively (Fig. 3). These numbers correspond to about 40% of the total pairwise interaction variance estimates (Fig. 2b), and greatly exceed expectations from background linkage effects^11^ (Supplementary Fig. 5). Like the detected additive QTL, the detected QTL-QTL interactions generally have very small effect sizes, with a median variance explained of 0.31%. The remainder of the interaction variance is likely due to many more pairs with even smaller effect sizes. Unlike the case for additive QTL, no large-effect QTL-QTL interactions were observed for these 20 traits. Whereas the largest additive QTL explained 26% of phenotypic variance, and 46 QTL had effect sizes greater than 5%, the largest QTL-QTL interaction explained only 3.3% of phenotypic variance (Fig. 4). Typical genetic architectures for traits in this study consist of a few large additive QTL and many small QTL and QTL-QTL interactions.

**Figure 4.**
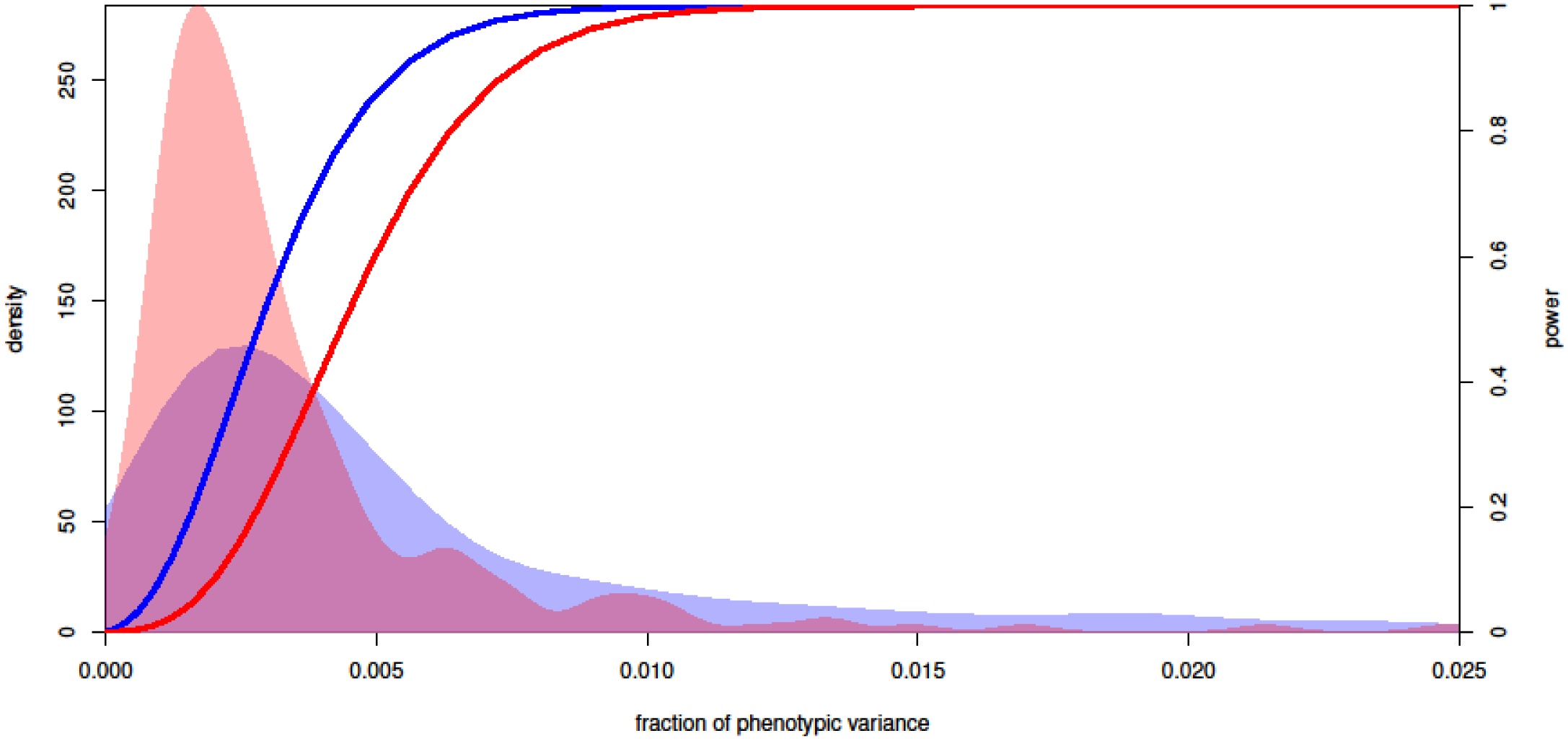
Distribution of genetic effects and power to detect them. A density plot of the fraction of phenotypic variance (X-axis) explained by individual significant QTL (blue area) and QTL-QTL interactions (red area) across all traits. The curves correspond to the statistical power at a genome-significance threshold (right Y-axis) for QTL (blue) and QTL-QTL interactions (red).

## Discussion

We have used a very large yeast cross with 4,390 segregants to study quantitative trait variation in greater detail. Across 20 traits, we find that additive genetic effects and pairwise genetic interactions explain 43.7% and 9.2% of phenotypic variance, respectively, in agreement with previous estimates based on a smaller dataset^12^. We detected a median of 42.5 significant additive QTL per trait. On average, these QTL captured 92% of the estimated additive heritability. Loci that explain at least 1% of phenotypic variance of loci typically spanned no more than 10 kb. We further estimate that roughly half of the pairwise interaction variance is contributed by interactions among significant additive QTL, and that nearly all of the interaction variance is contributed by interactions between significant additive QTL and the genome. Two-locus interactions in which neither locus has an additive effect are rare and do not contribute much to phenotypic variance. We detected about 13 QTL-QTL interactions per trait; these capture 3.2% of phenotypic variance or 40% of total pairwise interaction variance.

We previously discussed the factors that may lead to greater “missing”^13^ heritability in human genome-wide association studies (GWAS) than in a yeast cross^6^. These include greater genetic variation captured by population studies, differences in the allele frequency spectrum, larger mutational target size of the human genome, higher physiological complexity, and within-locus dominance effects. Here we have focused on better delineating the contributions of pairwise interactions to phenotypic variance. The larger cross enabled us to obtain an accurate genome-wide estimate of these contributions, and revealed that they are substantially smaller than those of additive effects for every trait examined. Further, few interactions arise from locus pairs without detectable additive effects. This is consistent with what has been observed in reverse-genetic screens with gene knockouts^14^. Although accurate estimates of the contributions of higher-order interactions require even larger sample sizes, the preliminary estimates obtained here (Supplementary Data 2) suggest that such interactions contribute less than pairwise ones. Theoretical results have been used to argue that the contributions of interactions to phenotypic variance in outbred populations are expected to be smaller than in a cross^1,2^. We note that a small contribution of genetic interactions to trait variance does not imply that interactions do not exist, that they are not important for understanding the complete genetic basis of specific traits, or that genes do not act epistatically at the molecular level^5,14^. Individual examples of QTL-QTL interactions, including some of large effect, have been detected for a broad range of traits in many species^5,15–17^. In studies that have estimated the contribution of pairwise interactions to trait variance, it is often within the range observed here ^18–21^. Our results further support the predominance of additive factors in explaining quantitative trait variance. They also suggest that interactions are most effectively detected by starting with the set of loci with additive effects. Combined with the recent observation of a small contribution of dominance to human trait variation^22^, this suggests that heritability not captured by genome-wide additive models arises primarily from additive effects of variants untagged by current genotyping technologies^23^.

## Methods

### Construction of segregant panel and sequencing libraries

The BYxRM segregants were constructed as described previously^6^. Before, we chose one segregant each from a panel of 1184 dissected tetrads, ultimately analyzing a panel of 1008 segregants. Here we added segregants from this panel of tetrads that were not previously genotyped to assemble a new panel of 4390 segregants. A Biomek FX liquid handling robot (Beckman Coulter) was used to re-array segregants that had not been previously genotyped to 1 ml of yeast peptone dextrose (YPD) in 2-ml deep-well 96-well plates (Thermo Scientific). Plates were sealed with Breathe-Easy gas-permeable membranes (Sigma-Aldrich), and yeast were grown 2 days at 30°C without shaking. DNA was extracted using 96-well DNeasy Blood & Tissue kits (Qiagen). DNA concentrations were determined using the Quant-iT dsDNA High-Sensitivity DNA quantification kit (Invitrogen) and the Bio-Tek Synergy 2 plate reader. DNA was diluted to 0.2 ng per microliter. Per sample, 5 µl of 0.2 ng per µl DNA was added to 4 µl of 5X Nextera HWM buffer (Illumina), 6 µl of water and 5 µl of 1/35 diluted Nextera enzyme. The transposition reaction was performed for 5 minutes at 55°C. Illumina sequencing adapters and custom indices were added by PCR directly after the tagmentation reaction without additional sample purification. 10 µl of fragmented DNA was combined with 0.5 µl each of 10 µM index primers (one of N701-N712 plus one of 96 custom indices), 5 µl of 10X Ex Taq buffer, 0.375 µl Ex Taq polymerase (Takara), 4 µl of 2.5 mM dNTPs, and 29.625 µl of water, and amplified with 20 cycles of PCR. 1152-plex libraries were run on two single end lanes of a rapid-run flow cell of a HiSeq 2500 (Illumina).

### Power calculations

We calculated statistical power (1-β) for sample sizes of 100, 1000, and 4000 segregants in R using the ‘power.t.test’ function^24^. Power was calculated over a range of effect sizes, where effect size was calculated as the percent phenotypic variance explained by a single QTL or QTL-QTL interaction. To correct for multiple testing genome-wide significance thresholds (α) of p<6.9x10^-4^ and p<2.5x10^-5^ were used for additive and interacting QTL, respectively. These thresholds were chosen based on a familywise error rate (FWER) < 5% for the additive scan and false discovery rate (FDR)<10% for the interaction scan. We note that we used a less stringent 10% FDR threshold for detecting individual QTL-QTL interactions to provide greater sensitivity to detect interaction effects and we expect that detected interactions are likely more upwardly biased than additive QTL effect sizes.

### Determining segregant genotypes

Fastq files for the 3,552 segregants sequenced for the present study were demultiplexed using fastq-multx^25^ and aligned to the SacCer3 version of the reference genome using bwa^26^. The 3,552 new segregants were sequenced with an average coverage of approximately 2X. The 1,056 previously sequenced segregants were realigned to SacCer3. BAM^27^ files for all 4,608 segregants were merged into one BAM file and variants were called as described previously. An additional filter was used to remove regions with strong mapping bias towards the reference genome^28^. Of 39,741 high confidence SNPs at which BY and RM differ, 28,220 unique SNPs were retained for downstream analysis. As described previously, a hidden Markov model was used to infer the segregant genotypes^6^. Segregants were removed if they had fewer than 25 or greater than 105 recombination breakpoints, fewer than 35,000 markers with genotype calls, or if the segregant genotype was correlated with another segregant with a Pearson correlation greater than 0.9. 4,390 segregants passed these filters and were used for mapping.

### Segregant phenotyping

All 4,390 segregants were phenotyped together, including the 1,056 previously characterized and sequenced segregants. Phenotyping was performed as described previously^6^. Briefly, segregants were pinned to agar plates from liquid stocks and then imaged for end-point growth at 48 hours. Colony radii were calculated using functions in the EBImage R package^29^. End-point growth measurements were filtered and normalized as previously described. Traits with larger difference between broad and narrow-sense heritabilities in our previous paper were prioritized here in order to focus on those traits more likely to have an appreciable contribution from genetic interactions. Therefore, the fraction of variance explained by genetic interactions could be biased upwards relative to all traits.

Segregant genotypes and phenotypes are available as Supplementary Data 5.

### Calculating variance components

Custom R code was used to estimate variance components and map additive QTL as well as QTL-QTL interactions. A repeated measures mixed model^7^ was used to estimate variance components. The model can be written as:

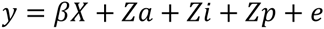

where *y* is a vector of length *m* that contains phenotypes for *n* segregants including replicate measurements such that *m = n * [number of replicates]*. β is a vector of estimated fixed effect coefficients. *X* is a matrix of fixed effects (here β is the overall mean, and *X* is a 1_*m*_ vector of ones unless otherwise specified). *Z* is an *m x n* incidence matrix that maps *m* total measures to *n* total segregants. *a* are the additive genetic effects, *i* are the pairwise genetic interaction effects and *p* are effects due to residual strain repeatability. The residual error is denoted by *e*. The distributions of these effects are assumed to be normal with mean zero and variance-covariance as follows:

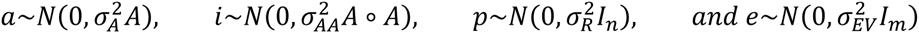

The variance structure of the phenotypes is 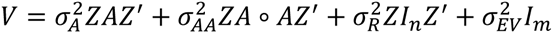. *A* is the additive relatedness matrix, the fraction of genome shared between pairs of segregants. *A* was calculated using the ‘A.mat’ function in the rrBLUP R package^30^. 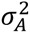 is the additive genetic variance captured by markers. *A*◦*A* is the Hadamard (entrywise) product of *A*, which can be interpreted as the fraction of pairs of markers shared between pairs of segregants. 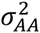 is the interaction genetic variance captured by all pairwise combinations of markers. *I*_*n*_ and *I*_*m*_ are *n x n* and *m x m* identity matrices, 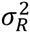 is the residual effect of strain not captured by the additive and interaction genetic variance terms, and 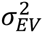 is the error variance. Variance components were estimated using AI-REML^31^ and custom R code. Standard errors of variance component estimates were calculated as the square root of the diagonal of the Fisher information matrix from the iteration at convergence of the AI-REML algorithm.

An additional term for three-way interactions, using the Hadamard cube of *A*, is included in a model in Supplemetary Data 2.

### Mapping additive QTL

Additive QTL were mapped using a forward stepwise procedure. For each chromosome and trait the above model was fit, replacing the term for polygenic additive effects with *a*_*loco*_ where 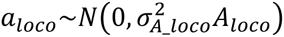. 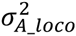 is the additive genetic variance from all chromosomes excluding the chromosome of interest, and *A*_*loco*_ is calculated as above, excluding markers from the target chromosome. The segregant best linear unbiased predictor (BLUP) residuals (*y*_*r*_ = *y* − *y*_*b*_) for each chromosome were calculated by subtracting the BLUPs for the effects of the rest of the genome and pairwise interactions from the phenotypes, where 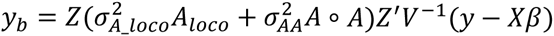 and 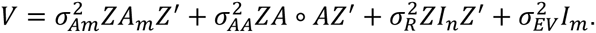. Replicate values per strain were averaged. These averaged BLUP residuals for each chromosome were then used as the starting point for scans for additive QTL on the chromosome of interest. Using BLUP residuals increases power to detect QTL by controlling for genetic contributions from the remainder of the genome^32^. We tested for linkage at each marker on the given chromosome by calculating *(-n(ln(1-r^2^)/2ln(10)))*, where *r* is the Pearson correlation coefficient between the segregant genotypes at the marker and segregant BLUP residuals for *n* segregants. FWER thresholds were determined from empirical null distributions determined by recomputing the linkage statistic chromosome wide from 1,000 permutations of BLUP residual phenotypes to strain assignments and recording the maximum value^33^. The most significant marker was extracted from each QTL significant at a 5% FWER threshold. These peak markers were added to the model as fixed effects and residuals were recomputed. Additional linkage scans were performed on these residuals (using 5% FWER thresholds that were recomputed after each round of QTL addition) until no additional significant QTL were detected on that chromosome. Confidence intervals were calculated as 1.5 LOD drop using the lodint function in R/QTL^34^.

### Mapping QTL-QTL interactions

We increased power and computational efficiency by searching for interactions using the segregant BLUP residuals from the additive polygenic model as phenotypes. Specifically, we calculated *y*_*r*_ for each trait as *y*_r_ = *y* − *y*_b_ where 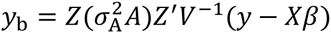 and 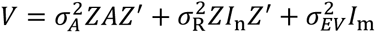. Replicate values per strain were averaged. For the full two-dimensional scan, LOD scores for interactions were computed for all pairs of markers as *(-n(ln(1-r^2^)/2ln(10)))*, where *n* is the number of segregants with phenotypes, and *r* is the Pearson correlation coefficient between the product of segregant genotypes at pairs of markers separated by at least 50 markers and the BLUP residuals. FDR at different LOD thresholds was calculated by dividing the average number of peaks obtained from 5 permutations of segregant identities by the number of peaks observed in the real data. We also tested for interactions between each locus with significant additive effects (identified as described in the preceding section) and the rest of the genome in the same manner as for the full two-dimensional scan. We refer to this as the marginal scan. FDR was calculated as above.

Results from the BLUP residual approach were compared to a simpler two locus interaction model from ‘scantwo’ in R/QTL^34^ that compares the likelihood ratio of a model that includes an interaction term to a model without this term. From the BLUP residual approach we detected 205 QTL-QTL in the full scan and 266 in the marginal scan. Using the same FDR procedure, 73 QTL-QTL were detected using R/QTL with the full two-dimensional scan and 112 were detected in the marginal scan. All of the R/QTL QTL-QTL interactions were also detected as statistically significant in our BLUP residual models.

### Fraction of variance captured by marker subsets

To estimate the fraction of additive variance captured by significant additive QTL, we fit the model *y* = *βX* + *Za* + *Zp* + *e*, where *a* was calculated from the relatedness of segregants only at the genome-wide significant QTL peak markers for the given trait (*A_QTL_*) such that 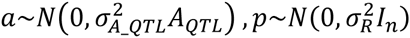 and 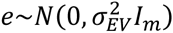, and compared it to the same model but with *a* calculated using the relatedness at all markers in the genome (*A*) as described above, such that 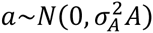.

We partitioned the interaction variance in a similar manner. Starting with *y* = *β* + *Za* + *Zi* + *Zp* + *e*, where 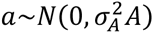 and 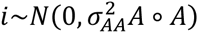, we replaced *A*◦*A* with various subsets of marker combinations. We fit a model with 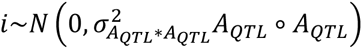 to capture the fraction of variance due to all pairwise interactions between significant additive QTL. We fit a model with 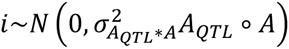 to capture the fraction of variance due to all pairwise interactions between significant additive QTL and the genome. We fit models with 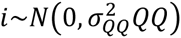, where 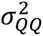 is the fraction of phenotypic variance captured by significant QTL-QTL interactions and *QQ* is the relatedness matrix calculated from an *n x q* matrix where *n* is the number of segregants and each column corresponds to the product of the genotypes at the peak markers for genome wide significant interacting QTL-QTL. The median fraction of interaction variance explained by significant QTL-QTL interactions was calculated as the median of 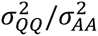 for the given trait.

To estimate the fraction of variance explained by non-specific background linkage effects, *N* markers or pairs of markers were chosen per trait, where *N* was the observed number of QTL or QTL-QTL for that trait. GRMs were calculated as above, but for the random marker subsets instead of QTL peak markers. To make this analysis more tractable, phenotype replicates were averaged for each strain and the repeatability term was excluded from the model. Variance components were estimated for each of the models listed above for 50 random draws of *N* markers for each trait. The median fraction of variance explained from these simulations is plotted in Supplementary Fig. 5.

The individual QTL and QTL-QTL interaction effect sizes shown in Fig. 4 were computed using the anova function in R with a trait specific multiple regression linear model with all the trait specific significant QTL peak markers and the product of QTL-QTL pair peak markers as fixed effects.

### Simulation of additive and pairwise interaction genetic architectures

We simulated phenotypes from a range of genetic architectures to test whether the mixed model will appropriately partition variance into additive and interaction components given our experimental design and our observed genotype data. Specifically, we simulated all combinations of either 0, 1, 5, 10, 50, or 500 QTL and / or QTL-QTL interactions. We set the broad-sense heritability (defined for these simulations as additive plus pairwise interaction variance) to 0.75 and varied the additive heritability to range from 0 to 0.75 in increments of 0.15 for all unique combinations of QTL and QTL-QTL interactions. QTL were given equal effects, but the sign of their effect was chosen at random. The positions of additive QTL were chosen randomly for each simulation. The positions of QTL-QTL interactions were chosen from the set of all combinations of additive QTL, but if the target number of QTL-QTL interactions was greater than the set of all combinations of additive QTL, then additional QTL-QTL interaction positions were chosen where neither position had a marginal additive effect. The summed effects of the additive loci were scaled to have the target additive variance and the summed effects of the interacting loci were scaled to have the target interaction variance and these were added to create vector *g*. Error variance was added from a normal distribution with mean 0 and *SD=(1-H2)/H2*var(g))*. Additive and interacting variance components were estimated with GRMs constructed from all the markers, as described above (Supplementary Data 1). We observed very large estimation errors in the case of architectures dominated by one very large effect interaction, but note that we did not observe such architectures for the traits studied here.

## Contributions

Experiments were designed by J.S.B. and L.K. Experiments were performed by J.S.B. and I.K. The genotyping protocol was developed by S.T. and I.K. Analyses were conducted by J.S.B. The manuscript was written by J.S.B. and L.K. and incorporates comments by M.S., F.W.A. and S.T.

## Acknowledgements

This work was supported by National Institutes of Health (NIH) grant R01 GM102308, a James S. McDonnell Centennial Fellowship, and the Howard Hughes Medical Institute (L.K.).

## Author Information

The authors declare no competing financial interests.

## Supplementary Figures

**Figure S1.**
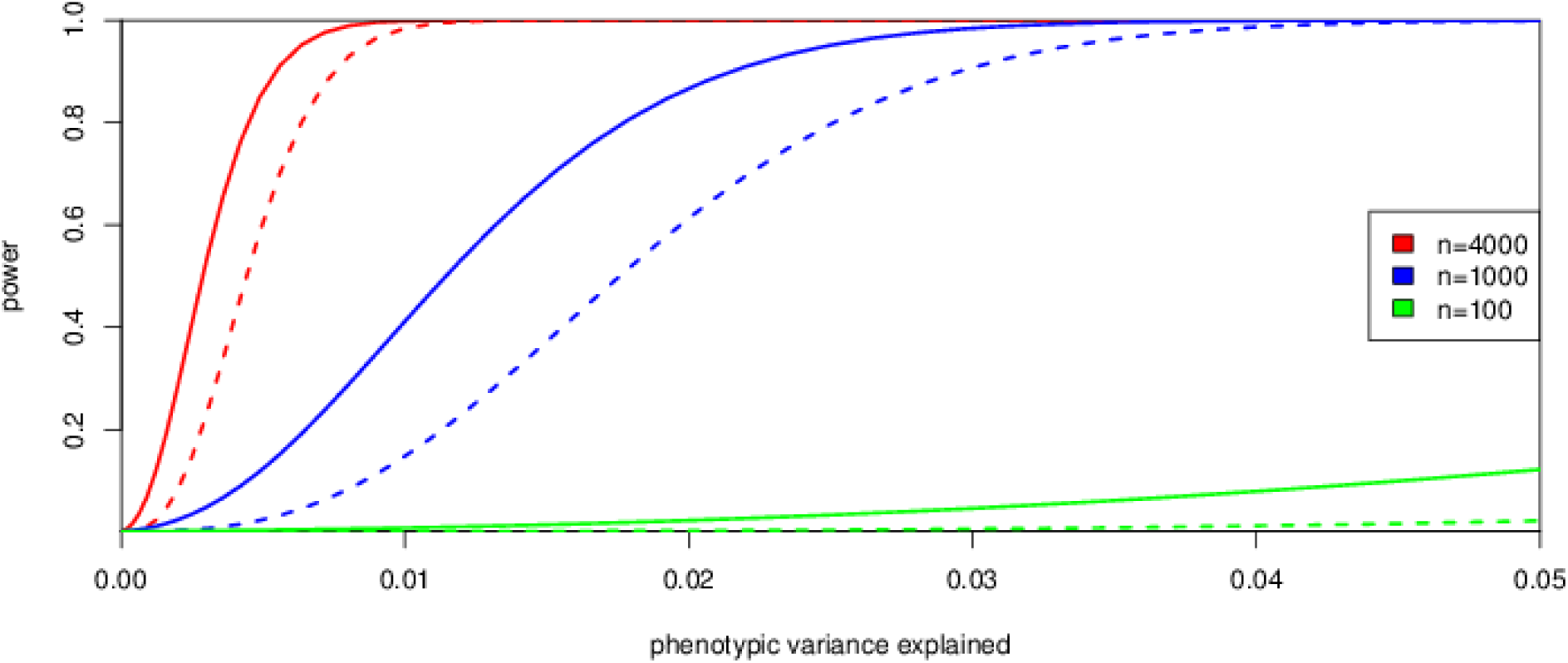
Curves illustrating statistical power are shown for mapping populations of 4000 (red), 1000 (blue), and 100 (green) segregants at a genome-wide significance threshold. The solid curves correspond to power for additive QTL and the dashed curve corresponds to power for QTL-QTL interactions.

**Figure S2.**
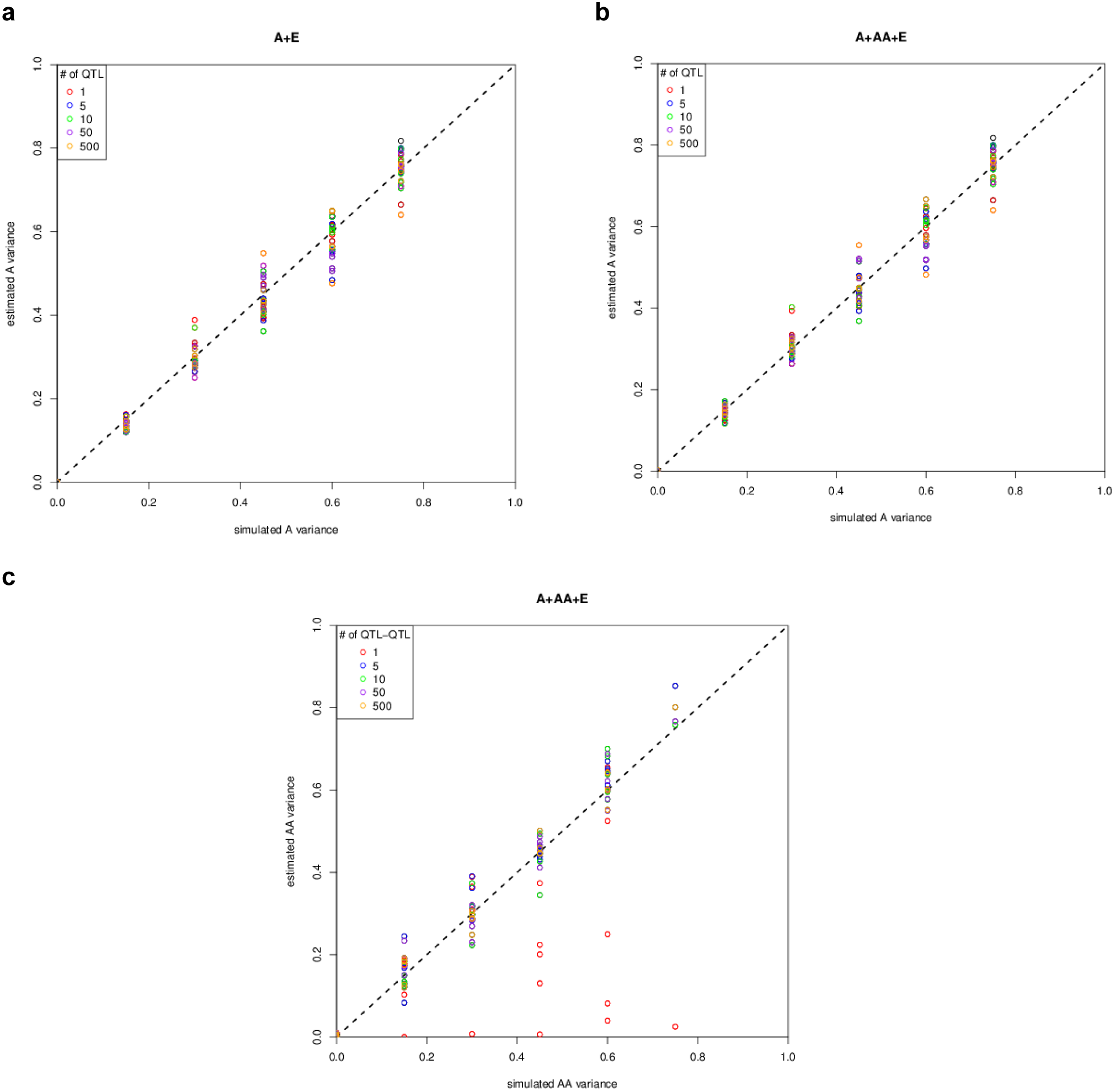
The amount of simulated variance (X-axis) is plotted against whole genome based variance component estimates (Y-axis). Each point represents a particular simulation with a specific genetic architecture. Colors indicate the number of QTL(a and b) or QTL-QTL interactions (c) simulated per simulated trait. a) Estimate of the additive variance component with A+E model. b) Estimate of the additive variance component with an A+AA+E model. c) Estimate of the interaction variance (AA) component with an A+AA+E model. Simulations include a range of QTL with QTL-QTL architectures ranging from 0 to 500 for each simulated trait, adding up to a total contribution of additive (a and b) or interaction variance (c) indicated on the X-axis. Simulations include interactions, even for the A+E model. The dotted black line represents (estimated = stimulated variance) and is shown as a visual guide.

**Figure S3.**
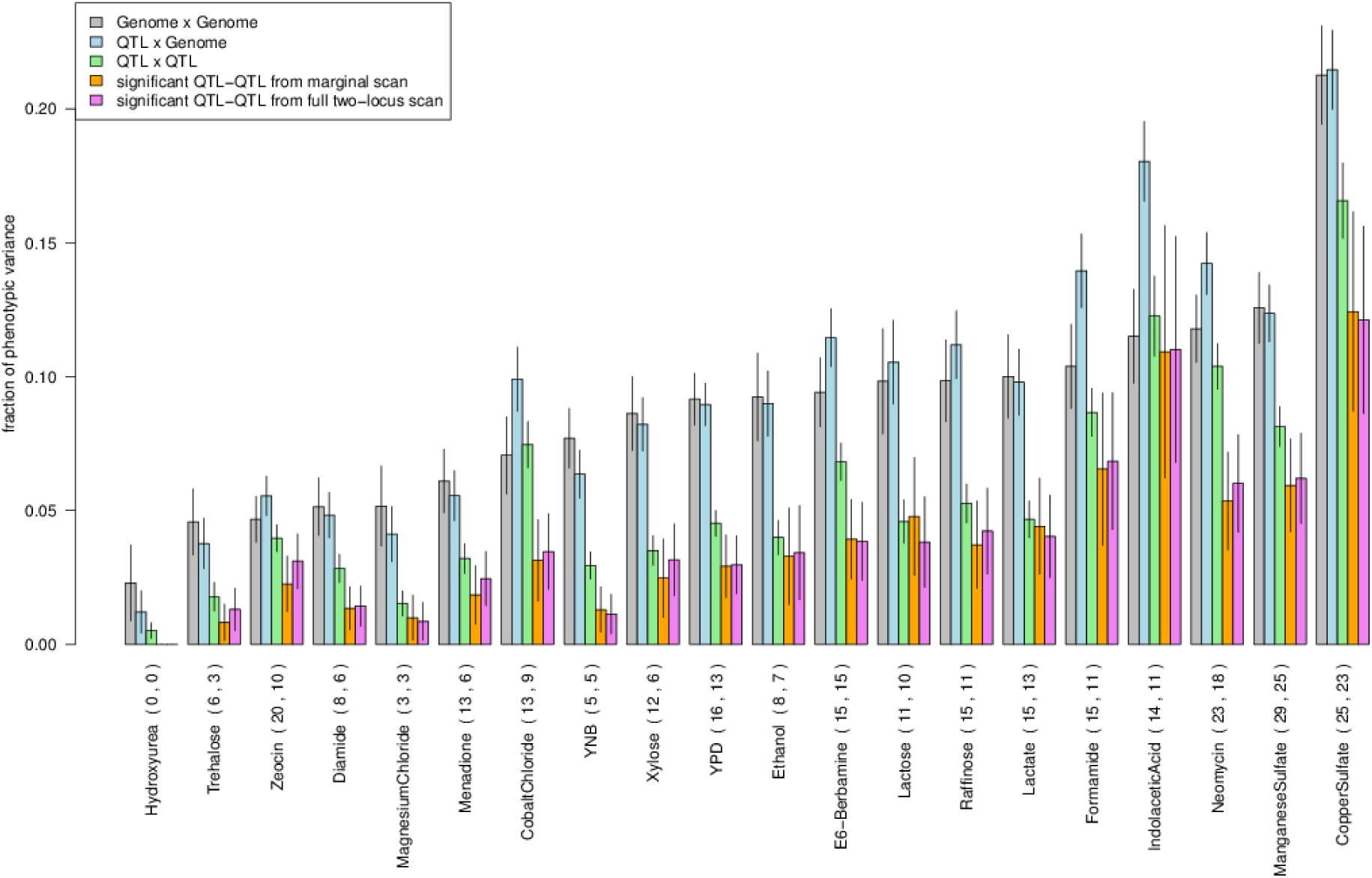
For each trait, juxtaposed barplots of the two-way interaction variance captured with all markers (genome x genome) (grey), significant additive QTL by all markers (QTL x genome) (blue), additive QTL by additive QTL (QTL x QTL) (green), significant QTL-QTL interactions detected from the marginal scan (orange), and significant QTL-QTL interactions from the exhaustive two-dimensional scan (purple) are shown. Error bars show +/− s.e.

**Figure S4.**
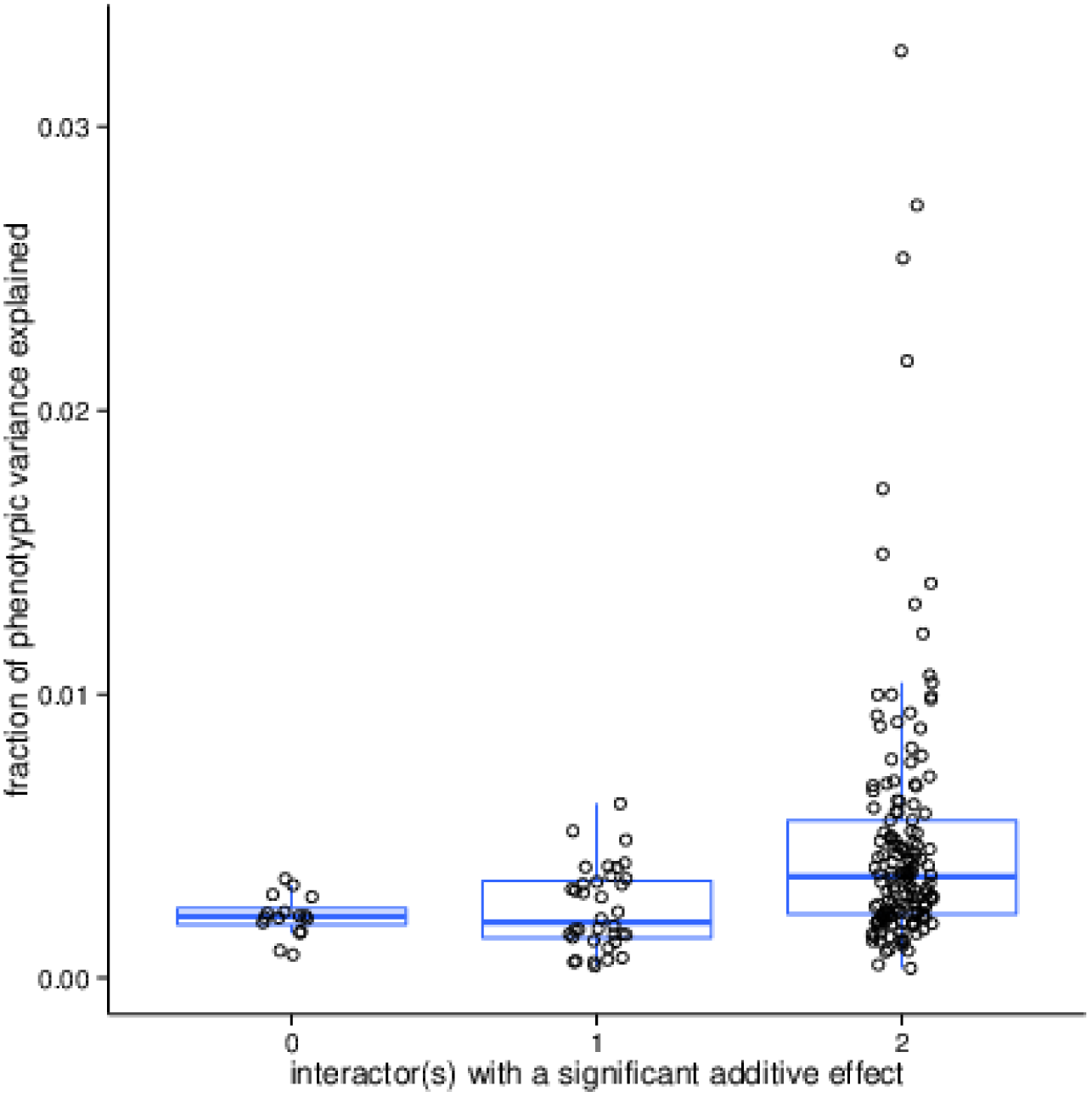
The fraction of phenotypic variance explained by individual significant QTL-QTL interactions from the exhaustive two-dimensional scan aggregated across all traits and grouped by whether 0, 1, or 2 of the interacting partners of the QTL-QTL interaction also have significant additive effects.

**Figure S5.**
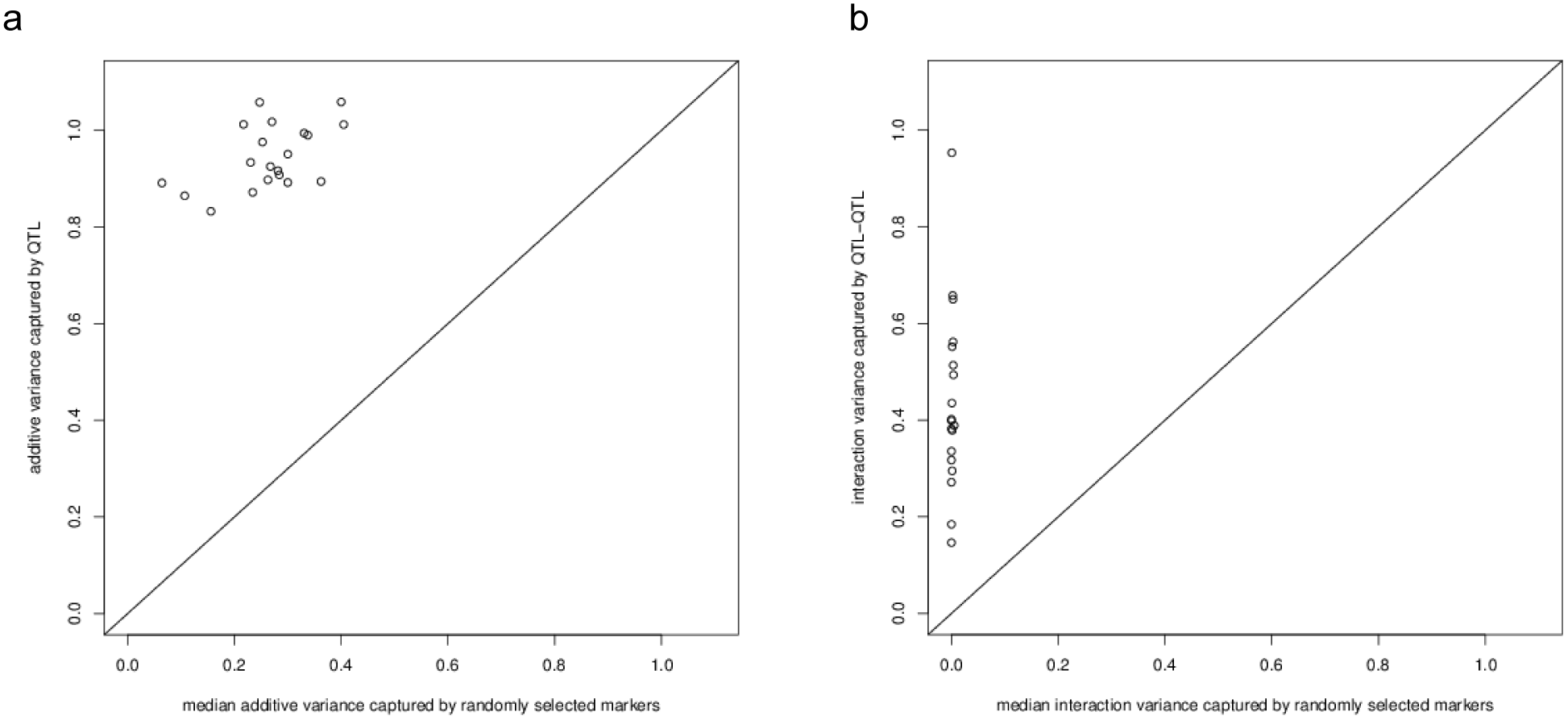
The median additive variance captured from N randomly selected markers (where N is the number of significant detected QTL for that trait) (X-axis) is plotted against the variance captured by QTL (Y-axis). The diagonal line represents (variance captured by detected QTL = variance captured due to background linkage effects) and is shown as a visual guide. B) The median interaction variance captured from N randomly selected pairs of markers (where N is the number of significant detected QTL-QTL pairs for that trait) (X-axis) is plotted against the variance captured by QTL-QTL pairs (Y-axis). The diagonal line represents (variance captured by detected QTL-QTL pairs = variance captured due to background linkage effects) and is shown as a visual guide.

**Supplementary Data 1. Simulation results**

Simulation results of different genetic architectures and additive and pairwise interaction variance component estimates. The numbers of additive QTL and QTL-QTL pairs, total simulated additive and interaction variances, and variance component estimates are indicated.

**Supplementary Data 2. Results of variance component analyses**

Results of variance component analyses for individual measurements for each trait.

**Supplementary Data 3. Detected additive QTL**

Positions, effect sizes, and variance explained by each QTL for each trait are listed.

**Supplementary Data 4. Detected QTL-QTL interactions**

Detected QTL-QTL interactions from the full and marginal interaction scan. Positions, effect sizes, and variance explained by each QTL-QTL interaction for each trait are listed.

**Supplementary Data 5. Segregant genotypes and phenotypes**

A zip file containing segregant genotypes and phenotypes as RData objects.

